# α-synuclein promotes neuronal dysfunction and death by disrupting the binding of ankyrin to ß-spectrin

**DOI:** 10.1101/2023.06.02.543481

**Authors:** Gali Maor, Ronald R. Dubreuil, Mel B. Feany

## Abstract

α-synuclein plays a key role in the pathogenesis of Parkinson’s disease and related disorders, but critical interacting partners and molecular mechanisms mediating neurotoxicity are incompletely understood. We show that α-synuclein binds directly to ß-spectrin. Using males and females in a *Drosophila* model of α-synuclein-related disorders we demonstrate that ß-spectrin is critical for α-synuclein neurotoxicity. Further, the ankyrin binding domain of ß-spectrin is required for α-synuclein binding and neurotoxicity. A key plasma membrane target of ankyrin, Na^+^/K^+^ ATPase, is mislocalized when human α-synuclein is expressed in *Drosophila*. Accordingly, membrane potential is depolarized in α-synuclein transgenic fly brains. We examine the same pathway in human neurons and find that Parkinson’s disease patient-derived neurons with a triplication of the α-synuclein locus show disruption of the spectrin cytoskeleton, mislocalization of ankyrin and Na^+^/K^+^ ATPase, and membrane potential depolarization. Our findings define a specific molecular mechanism by which elevated levels of α-synuclein in Parkinson’s disease and related α-synucleinopathies leads to neuronal dysfunction and death.

**Significance Statement:** The small synaptic vesicle associate protein α-synuclein plays a critical role in the pathogenesis of Parkinson’s disease and related disorders, but the disease-relevant binding partners of α-synuclein and proximate pathways critical for neurotoxicity require further definition. We show that α-synuclein binds directly to ß-spectrin, a key cytoskeletal protein required for localization of plasma membrane proteins and maintenance of neuronal viability. Binding of α-synuclein to ß-spectrin alters the organization of the spectrin-ankyrin complex, which is critical for localization and function of integral membrane proteins, including Na^+^/K^+^ ATPase. These finding outline a previously undescribed mechanism of α-synuclein neurotoxicity and thus suggest potential new therapeutic approaches in Parkinson’s disease and related disorders.

## Introduction

Parkinson’s disease is the most common neurodegenerative movement disorder, affecting 1% of individuals at age 65 (Van Den Eeden et al., 2003; de Lau and Breteler, 2006; Yang et al., 2019). There are currently no effective disease-modifying therapies for Parkinson’s disease and related disorders, emphasizing the importance of delineating the underlying pathobiology so that rational therapeutics can be developed. Genetics and neuropathology have converged to implicate strongly the small, 140 amino acid, protein α-synuclein in the pathogenesis of Parkinson’s disease and related diseases. Point mutations, duplications and triplications of the α-synuclein locus cause autosomal dominant, highly penetrant familial forms of Parkinson’s disease (Polymeropoulos et al., 1997; Krüger et al., 1998; Zarranz et al., 2004; Book et al., 2018). In both rare forms of genetic Parkinson’s disease linked to α-synuclein mutations and in more common forms of the disorder α-synuclein is deposited into a variety of intracellular protein aggregates. Aggregation of α-synuclein is also seen in the related neurodegenerative disorders dementia with Lewy bodies and multiple system atrophy. These diseases are collectively termed α-synucleinopathies.

The implication of elevated α-synuclein levels as a cause of Parkinson’s disease has led to the development of animal models based on increased expression of human α-synuclein. In monkeys, rats, mice, fish, flies and worms expression of human α-synuclein can mimic key features of Parkinson’s disease including progressive locomotor dysfunction, degeneration of dopaminergic and non-dopaminergic neurons, and α-synuclein aggregation (Feany and Bender, 2000; Masliah et al., 2000; Kirik et al., 2002, 2003; Lee et al., 2002; Lakso et al., 2003; O’Donnell et al., 2014). Since flies and worms do not normally express α-synuclein, toxicity plausibly represents gain of function of the human protein. Using these animal α-synucleinopathy models, a variety of processes including phosphorylation and aggregation of α-synuclein (Chen and Feany, 2005; Lo Bianco et al., 2008; Chen et al., 2009; Kuwahara et al., 2012), mitochondrial dysfunction (Martin et al., 2006; Ordonez et al., 2018; Sarkar et al., 2020; Portz and Lee, 2021) and altered proteostasis (Auluck et al., 2002; Colla et al., 2012; Yan et al., 2019; Karim et al., 2020; Sarkar et al., 2021) have been implicated in α-synuclein neurotoxicity. However, the proximal mechanisms linking α-synuclein to downstream mediators have remained unclear. We now show that α-synuclein binds directly to ß-spectrin in vitro and mediates neurotoxicity in vivo by disrupting ankyrin- and spectrin-dependent localization and function of the plasma membrane Na^+^/K^+^ ATPase.

## Materials and Methods

### *Drosophila* stocks and Genetics

All fly crosses and aging were performed at 25°C. Equal numbers of adult male and female flies were analyzed at 10 days post-eclosion except as otherwise indicated in the figure legends. The *QUAS-*α*-synuclein wild type* transgenic flies been reported previously (Ordonez et al., 2018). N-terminally myc-tagged β*-spectrin wild type* (β*-spectrin^KW3A^*), β*-spectrin^Δank^* (β*-spectrin*^α*13*^) and β*-spectrin*^Δ*PH*^ transgenes were expressed under the control of the *Drosophila* ubiquitin promotor. These β-spectrin transgenic flies have been described in detail previously (Das et al., 2006). Expression of α-synuclein was directed to neurons using the pan-neuronal driver *nSyb-QF2*. GAL4-mediated expression was controlled by the pan-neuronal *nSyb-GAL4* driver, from the Bloomington *Drosophila* Stock Center. The *nSyb-QF2* line was obtained from C. Potter.

### Behavioral analysis

The climbing assay was performed as described in detail (Ordonez et al., 2018). Briefly, ten flies were placed in individual vials and a total of six vials assayed per genotype, for a total of 60 flies per genotype. Flies were tapped down gently to the bottom of the vial, and the number of flies climbing above 5 cm within 10 seconds was recorded. Results are reported as the mean plus and minus the standard error of the mean.

### iPS cells and neuronal differentiation

NGN2-induced pluripotent stem cells from a female donor were obtained from Brigham and Women’s iPSC Neurohub and maintained as feeder-free cells in a defined, serum-free media (mTeSR, Stemcell Technologies). For neuronal induction, cells were dissociated with Accutase (Stemcell Technologies) and plated in mTeSR supplemented with 10 μM ROCK inhibitor Y-27632 and 2 μg/mL doxycycline on a Matrigel coated 6-well plate. On day one of the differentiation, culture media was changed to DMEM/F12 supplemented with N2 (Gibco), B27 (Gibco), non-essential amino acids, GlutaMAX, 5 µg/ml puromycin and 2 µg/ml doxycycline. On day four of differentiation, media was changed to Neurobasal media (Gibco) supplemented with B27 (Life Technologies), 10 ng/µl BDNF, CNTF and GDNF, 10 µM ROCKi, 5 µg/ml puromycin and 2 µg/ml doxycycline. Medium was changed every three days. For the imaging studies reported at least three independent differentiations of triplication and isogenic control neurons plated in parallel were performed and analyzed.

### Histology, immunohistochemistry and immunofluorescence

For examination of the adult fly brain, animals were fixed in formalin and embedded in paraffin. 4 μm serial frontal sections were prepared through the entire brain and placed on a single glass slide. Hematoxylin staining was performed on paraffin sections to assess total neuronal density. Neurons differentiated from iPS cells were fixed in 4% paraformaldehyde before proceeding to immunostaining.

For immunostaining of paraffin sections, slides were processed through xylene, ethanol, and into water. Antigen retrieval by boiling in sodium citrate, pH 6.0, was performed prior to blocking. In some studies, whole mount *Drosophila* brain preparations were alternatively used. Blocking was performed in PBS containing 0.3% Triton X-100 and 2% milk for 1 hour and followed by incubation with appropriate primary antibodies overnight. Primary antibodies used were: anti-tyrosine hydroxylase (Immunostar) at 1:500; anti-α-synuclein (5G4, Millipore) at 1:1,000,000, anti-β-spectrin (Dubreuil laboratory) at 1:1000; anti-nrv1 (Nrv5F7, Developmental Studies Hybridoma Bank) at 1:100; anti-ankyrin/ankyrin B (N105/17, NeuroMab) at 1:200, anti-βII-spectrin (BD Biosciences) at 1:500; anti-ATP1B1 (Abcam), anti-ßIV-spectrin (polyclonal antibody from Dr. M. Rasband) at 1:500, anti-ankyrin G (N106/36, NeuroMab) at 1:200, anti-tubulin β3 (Biolegend) at 1:500, and anti-MAP2 (Abcam) at 1:5,000. For immunohistochemistry, biotin-conjugated secondary antibodies (1:200, SouthernBiotech) and avidin-biotin-peroxidase complex (Vectastain Elite, Vector Laboratories) staining was performed using DAB (Vector Laboratories) as a chromagen. For immunofluorescence studies, appropriate Alexa Fluor conjugated secondary antibodies (Alexa 488, Alexa 555 or Alexa 647, 1:200, Invitrogen) were used.

For quantification of TH-positive neurons and aggregated α-synuclein in *Drosophila* brains, an entire tissue cross section of the anterior medulla was imaged. One image per fly and a total of 6 flies per genotype were used for quantification. The number of TH-positive cells detected by immunohistochemistry or α-synuclein aggregates detected with immunofluorescence in each image was counted and results were expressed per unit area.

### Cloning, protein purification and precipitation

Wild type *Drosophila* and human spectrins were cloned into a histidine tagged vector (2BC-T cloning vector, Addgene # 31070) using ligation independent cloning to obtain C-terminally tagged proteins. Briefly, the vector was linearized using HpaI digestion. Inserts encoding fly and human α- and β-spectrins were PCR-amplified using appropriate primers. Following amplification, constructs were annealed at room temperature for 5 minutes. Mutant *Drosophila* and human β-spectrins (β-spec^Δank^) were produced by VectorBuilder (https://en.vectorbuilder.com/). In both mutants, the sequence of the 15th β-spectrin repeat was replaced with the sequence of the 12th α-spectrin repeat, as in the β-spec^Δank^ transgenic flies (Das et al., 2006).

For isolation of recombinant protein, overnight cultures of transformed BL21□cells were induced with 1LmM IPTG at 37°C for 4□hours and harvested by centrifugation. Proteins were eluted using Ni-NTA spin columns according to the manufacturer’s protocol (Qiagen). GST-α synuclein fusion protein tagged at the N-terminus was obtained from Sigma Aldrich. 200 µg of GST-α synuclein was incubated with the equivalent amount of purified His-spectrin at 4°C overnight in the presence of glutathione sepharose beads. The pull-downs were washed 4 times with lysis buffer and finally resolved by boiling in 2X Laemmli sample buffer (63 mM Tris–HCl pH 6.8, 10% glycerol, 2% SDS, 0.0025% bromophenol blue) at 95°C for five minutes. 200 µg of the proteins was incubated with equivalent amount of GST-α synuclein fusion protein at 4°C overnight in the present of Ni-NTA beads (Invitrogen). The pull-downs were washed 4 times with lysis buffer and boiled in 2X Laemmli sample buffer for five minutes.

### Immunoprecipitation

To assess α-synuclein and β-spectrin interaction in vivo, immunoprecipitation was performed. 10 fly heads or differentiated neurons from a confluent 10 cm culture plate were homogenized in non-denaturing lysis buffer (20 mM Tris HCl pH 8, 137 mM NaCl, 1% Triton X-100, 2 mM EDTA) and centrifuged at 12,000 rpm to pellet debris. The supernatant was incubated with the H3C monoclonal α-synuclein antibody with rotation for 12 hours at 4°C. Protein-G Sepharose beads (GE Healthcare) were blocked in 0.1% BSA for 1 hour at room temperature, washed and added to cell lysates for incubation with rotation for 4 hours at 4°C. The precipitated material was then washed 4 times in lysis buffer, resuspended with SDS loading buffer and subjected to immunoblotting.

### Western blots

*Drosophila* heads were homogenized in 2X Laemmli sample buffer. All samples were boiled for 10 minutes, briefly centrifuged and subjected to SDS-PAGE using 10% gels (Bio-Rad). Proteins were transferred to nitrocellulose membranes (Bio-Rad), blocked in 2% milk in PBS with 0.05% Tween-20, and immunoblotted with primary antibodies. Primary antibodies used were anti-α-synuclein (H3C, Developmental Studies Hybridoma Bank) at 1:500,000; anti-c-Myc (9E10, Developmental Studies Hybridoma Bank) at 1:1,000; anti-6XHis (N144/14, NeuroMab) at 1:500; anti-GAPDH (Invitrogen) at 1:1000. The appropriate IRDye fluorescence secondary antibody (1:10,000, LICOR) was applied. Images were taken using a LICOR Odyssey DLx imaging system (LICOR). Blots were repeated at least three times, and a representative blot shown.

### Fluorescence microscopy

Confocal images were taken on a Zeiss LSM-800 confocal microscope with Airyscan. For the evaluation of βII-spectrin cytoskeleton, the axon initial segment, and ankyrin-B and Na^+^/K^+^ ATPase localization in neurons differentiated from iPSC three independent differentiations of triplication and isogenic control neurons plated in parallel were performed. For each parallel set of isogenic control and triplication neurons 3 coverslips were analyzed with a total of approximately 1,000 neurons analyzed for each for control and triplication cells. Imaged regions were selected based on optimal cell density to allow visualization of the soma, axon and dendrites of individual neurons, and on consistent immunostaining. All cells within an imaged field were analyzed and 1) the percentage of cells with disrupted βII-spectrin cytoskeleton, 2) the staining pattern of ankyrin-G and βIV-spectrin, and 3) localization of ankyrin-B and Na^+^/K^+^ ATPase was assessed. For quantification of ankyrin-B and Na^+^/K^+^ ATPase localization, the fluorescence intensity of staining in the plasma membrane as well as in the cytoplasm was measured in the soma. Plasma membrane staining was derived by subtraction of the cytoplasmic staining from total staining of the cells.

#### Plasma membrane polarization

The voltage sensitive fluorescent dye bis-(1,3-dibutylbarbituric acid) trimethine oxonol (DiBAC_4_(3)) was used in *Drosophila* brains and cultured neurons. Relative fluorescent intensities were recorded, higher intensities indicated depolarization of the plasma membrane. Brains from ten-day old flies were dissected in Schneider’s media and incubated with 4 µM DiBAC4(3) for 20 minutes at 25°C. Brains were then mounted and confocal microscopy performed immediately with quantification of average pixel intensity from two-dimensional projections of confocal z-stacks representing the entire brain using ImageJ. Imaging and analysis for all genotypes were done at the same confocal settings, including laser intensity and z-stack thickness. Results represent the average of 6 flies per genotype. Cultured cells were incubated in 200 nM DiBAC_4_(3) in DMEM/F12 for 20 minutes in 37°C incubator at 21 DIV. Cells were imaged without washing. Results represent an average of 100 cells per cell line. Imaging and analysis for both cell lines were done at the same confocal settings.

### Stimulated emission depletion (STED) microscopy

To analyze the spectrin cytoskeleton and cellular localization of ankyrin and Na^+^/K^+^ ATPase in *Drosophila* brains from 10-day-old flies were dissected and fluorescently labeled using standard protocols and mounted using Prolong Diamond antifade mounting medium (Invitrogen). Imaging of Kenyon cells was performed using a STED instrument mounted on a Leica SP8 confocal microscope. Fluorophores were excited at 488 or 550 nm from a white light laser and depletion of the signal was done at 592 and 660 nm, respectively. A time gate was used to reject photons with a lifetime outside a 1.3 and 6 ns time window. To achieve optimal resolution, pixel size was matched to be 25 nm, with averaging of 4 images. For quantification of spectrin cytoskeletal disruption and membrane and cytosolic ankyrin and Na^+^/K^+^ ATPase, a well-stained optical section from the midportion of the mushroom body Kenyon cell layer representing the entire thickness of the cortex and including approximately 30 cells was imaged. To quantify spectrin cytoskeletal disruption the number of cells with focal or multifocal discontinuities of the subplasmalemmal spectrin network was counted in each image. For analysis of ankyrin and Na^+^/K^+^ ATPase, the intensity from each cell in the image was measured. A total of 6 animals per genotype were assessed. For colocalization studies, the Pearson coefficient was calculated from 4 images per animal, each image containing 20 cells.

### Statistical analysis

Non-parametric statistical tests were used for all comparisons. Details regarding statistical tests, biological sample size (n) and p value are present in figure legends. All data are represented as mean ± SEM. SEM represents variance within a group. Data were collected and processed side by side in randomized order for all experiments. Unpaired, two-tailed t tests were used for comparison between two groups, with p < 0.05 considered significant. For all comparisons involving multiple variables, one-way or two-way ANOVA was performed followed by Bonferonni tests for multiple comparison using p < 0.05 for significance. All statistical analyses were preformed using GraphPad Prism.

## Results

We have previously described a *Drosophila* model of Parkinson’s disease and related α-synucleinopathies based on expression of wild type human α-synuclein in a pan-neuronal pattern. Our model recapitulates key features of the human disorder, including progressive locomotor dysfunction, age-dependent neurodegeneration and α-synuclein aggregation (Feany and Bender, 2000; Ordonez et al., 2018). In prior work we further observed that α-synuclein neurotoxicity depended on the levels of α-spectrin (Ordonez et al., 2018). Spectrins are highly conserved tetrameric cytoskeletal proteins consisting of two α and two ß subunits. *Drosophila* is a favorable model system for the study of spectrin function because spectrins are encoded by just three genes in flies: one α*-spectrin* gene, one *ß-spectrin* gene and one *ß_H_-spectrin* gene. We previously found that increased expression of α-spectrin could rescue downstream toxicity of α-synuclein, including mitochondrial dysfunction and neuronal death (Ordonez et al., 2018). However, other studies have suggested that ß-spectrin, as well as α-spectrin, can interact with α-synuclein (Leverenz et al., 2007; McFarland et al., 2008; Lee et al., 2012; Chung et al., 2017). We therefore expressed ß-spectrin together with α-synuclein in our fly model to determine if ß-spectrin can modulate α-synuclein neurotoxicity in vivo. We found, as reported previously, that α-synuclein transgenic flies displayed impaired locomotor function as assessed by the climbing assay. Elevated expression of transgenic ß-spectrin ameliorated locomotor dysfunction in α-synuclein transgenic flies (Fig. 1*A*). Similarly, degeneration as assessed by total numbers of cortical cells (Fig. 1*B,C*) and numbers of tyrosine hydroxylase-positive dopamine neurons (Fig. 1*D*, arrows, *E*) was also rescued by elevated expression of ß-spectrin. Rescue did not simply reflect reduced levels of α-synuclein since western blotting revealed equivalent levels of α-synuclein when ß-spectrin was increased in expression (Extended Data Fig. 1-1*A,B*).

**Figure 1.**
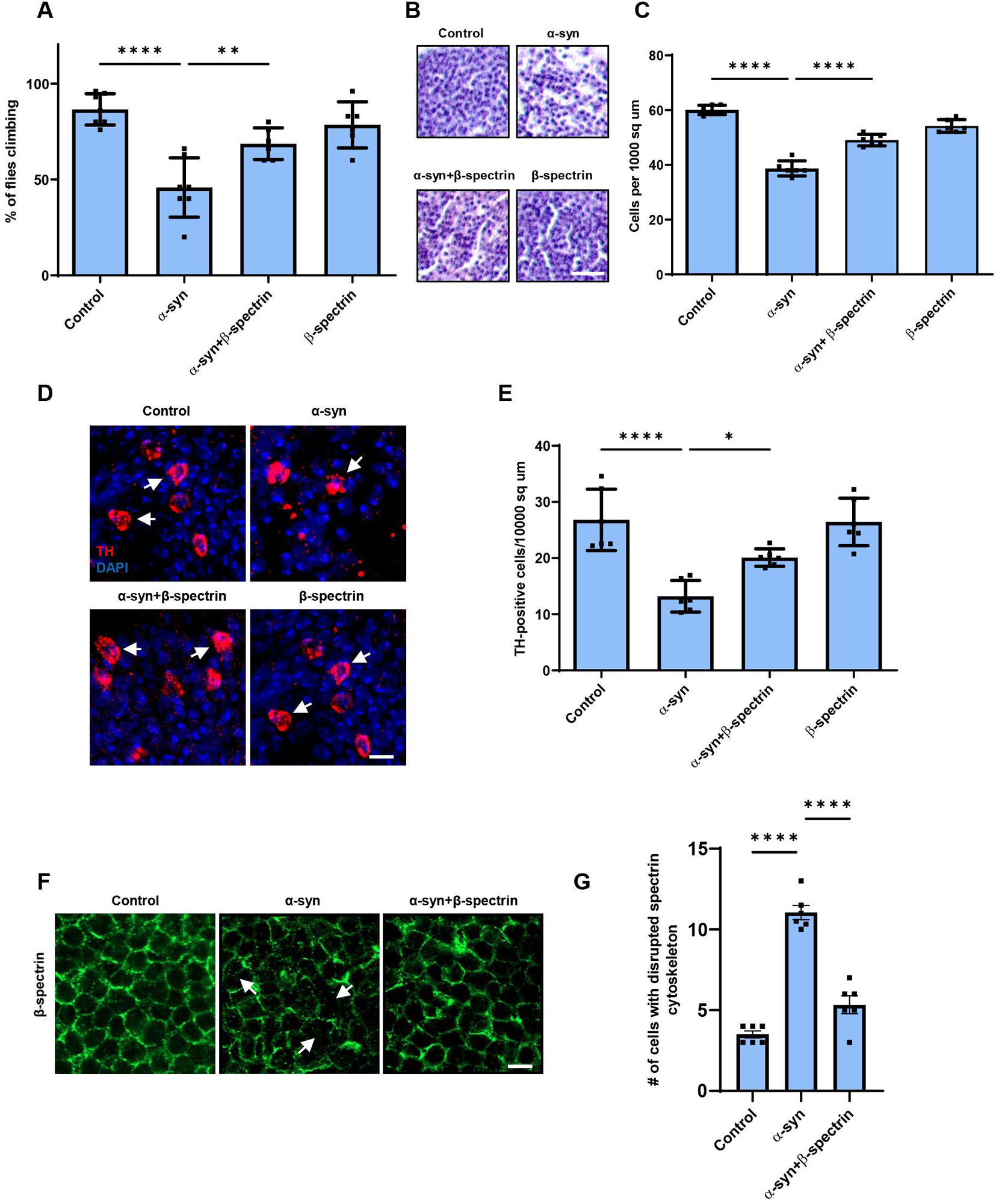
Elevated expression of β-spectrin rescues α-synuclein neurotoxicity. *A,* Climbing activity in control and human α-synuclein transgenic *Drosophila* with and without elevated expression of β-spectrin. n = minimum of 60 flies per genotype (six biological replicates of 10 flies each). *B,C*, Brain degeneration assayed by hematoxylin staining in the anterior medulla of control and α-synuclein transgenic *Drosophila* with and without elevated expression of β-spectrin; n=6. *D,E*, Immunostaining for tyrosine hydroxylase in the anterior medulla of control and α-synuclein transgenic *Drosophila* with and without elevated expression of β-spectrin. n=6. *F,G*, The normal spectrin subplasmalemmal network is disrupted by expression of human α-synuclein. Arrows indicate disruptions in the spectrin network. n=6. All flies are 10 days old. Controls are *nSyb-QF2, nSyb-GAL4/+.* The scale bars represent 10 µm (*B*) and 5 µm (*D*) and (*F)*. Data are presented as mean ± SEM; P values determined with one-way ANOVA with Bonferroni post hoc test. See Extended Data Fig. 1-1.

The spectrin cytoskeleton is anchored to the cytoplasmic face of the plasma membrane and is believed to form a network that contributes to maintaining the structure and shape of the cell (Dubreuil, 2006; Morrow and Stankewich, 2021; Teliska and Rasband, 2021). Immunostaining revealed regular subplasmlemmal ß-spectrin staining in the brains of control flies (Fig. 1*F*) (Das et al., 2008). In contrast, staining was irregular and disrupted in the brains of α-synuclein transgenic flies (Fig. 1*F*, arrows, *G*), similar to the pattern seen with perturbed ß-spectrin function (Das et al., 2008). Elevated expression of β-spectrin partially restored the ß-spectrin staining pattern in flies expressing α-synuclein (Fig. 1*F,G*).

We next examined the functional consequences of lowering levels of β-spectrin in α-synuclein transgenic flies. Animals with no β-spectrin die as late embryos or early larvae (Dubreuil et al., 2000; Das et al., 2006). We therefore used flies with a complete loss of function β-spectrin mutation (β*-spec^em21^*) combined with a transgene expressing an engineered mutant of β-spectrin with three amino acid substitutions in the actin binding domain (β-spectrin^Kpn+3^, Dubreuil, unpublished), which accumulates at significantly reduced levels (Extended Data Fig. 1-1) but rescues lethality of β-spectrin loss of function. We therefore used hemizygous male β*-spec^em21^* flies rescued to adulthood by one copy of the β-spectrin^Kpn+3^ transgene in our β-spectrin knockdown experiments, termed β-spectrin^KD^ here for simplicity (Fig. 2). α-synuclein transgenic flies with loss of β-spectrin function exhibited a significant enhancement of locomotor dysfunction (Fig. 2*A*). Similarly, degeneration as monitored by total numbers of cortical cells (Fig. 2*B,C*) or tyrosine hydroxylase-positive neurons (Fig. 2*D*, arrows, *E*) worsened with reduced β-spectrin expression.

**Figure 2.**
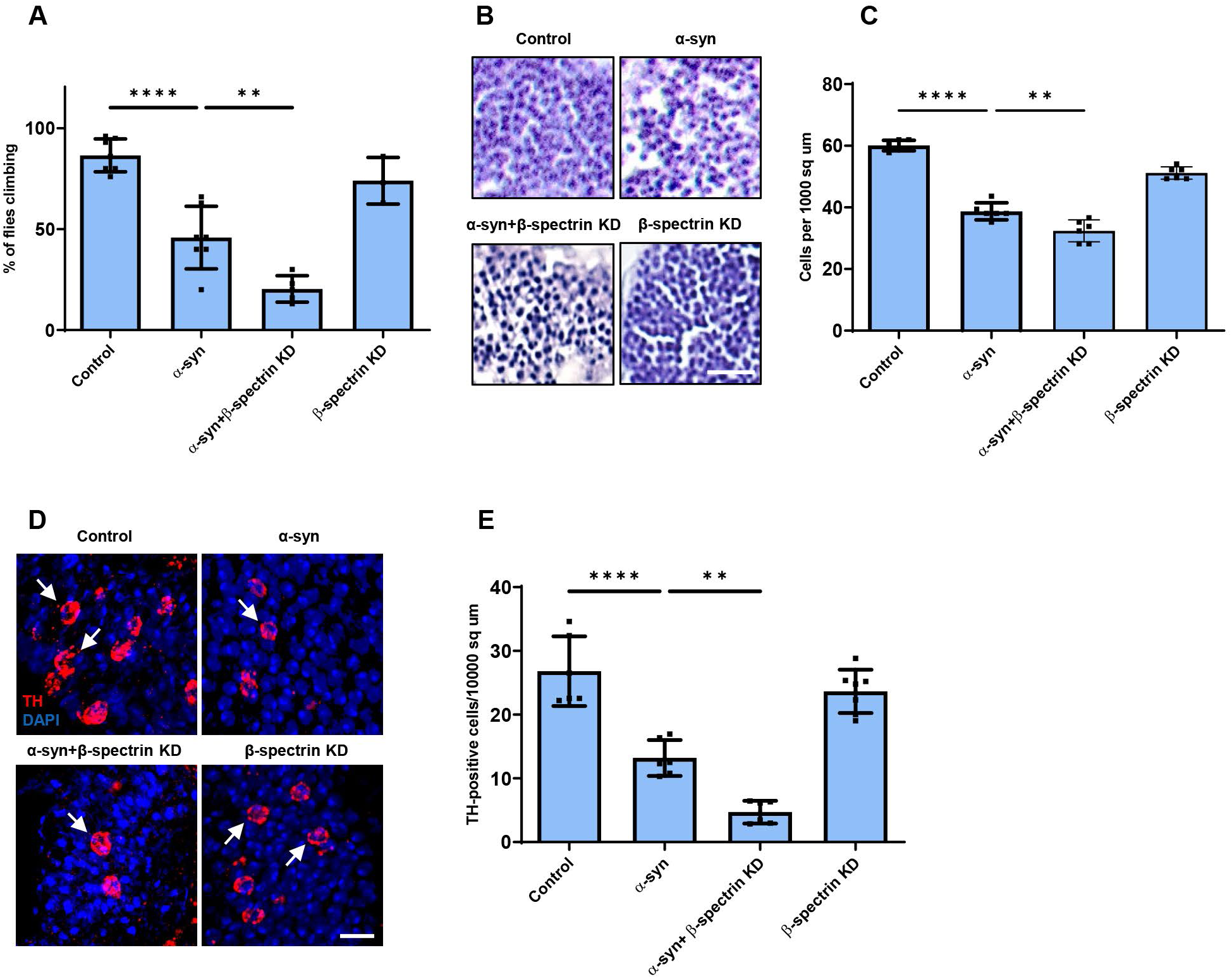
Reduced expression of β-spectrin enhances α-synuclein neurotoxicity. *A*, Climbing activity in control and α-synuclein transgenic *Drosophila* with and without reduced expression of β-spectrin. n = minimum of 60 flies per genotype (six biological replicates of 10 flies each). *B,C*, Brain degeneration assayed by hematoxylin staining in the anterior medulla of control and α-synuclein transgenic *Drosophila* with and without reduced expression of β-spectrin; n=6. *D,E*, Immunostaining for tyrosine hydroxylase in the anterior medulla of control and α-synuclein transgenic *Drosophila* with and without expression of β-spectrin. n=6. All flies are 10 days old. β-spectrin^KD^ flies are hemizygous for the complete loss of function mutation β*-spec^em21^*, rescued to viability with transgenic expression of the β*-spec^Kpn+3^* variant, which accumulates at significantly reduced levels. Controls are *nSyb-QF2, nSyb-GAL4/+.* The scale bars represent 10 µm (*B*) and 5 µm (*D*). Data are presented as mean ± SEM; P values determined with one-way ANOVA with Bonferroni post hoc test. See Extended Data Fig. 1-1.

To determine if α-synuclein interacts directly with spectrin, we expressed histidine-tagged α- and β-spectrin proteins in bacteria. Each spectrin was purified, incubated with purified, GST-tagged human α-synuclein and the samples precipitated with Ni-NTA or glutathione Sepharose beads. *Drosophila* β-spectrin bound to α-synuclein in vitro (Fig. 3*A*), which we verified in vivo by immunoprecipitation (Fig. 3*E*). We also determined that human βII-spectrin (Fig. 3*C*) bound to α-synuclein in vitro, as seen by both GST and Ni-NTA precipitation. We began by assessing human βII-spectrin because of the five mammalian β-spectrins, βI, βII, βIII, βIV and βV (Lorenzo, 2020), the βII isoform is abundant in the nervous system and is most similar to the conventional *Drosophila* β-spectrin. In contrast to the β-spectrins, interactions between *Drosophila* α-spectrin or human αII-spectrin and α-synuclein were not detected in vitro (Fig. 3*B,D*). Of the two mammalian α-spectrins, αII-spectrin is expressed in brain, while αI-spectrin is predominantly found in red blood cells (Lorenzo, 2020).

**Figure 3.**
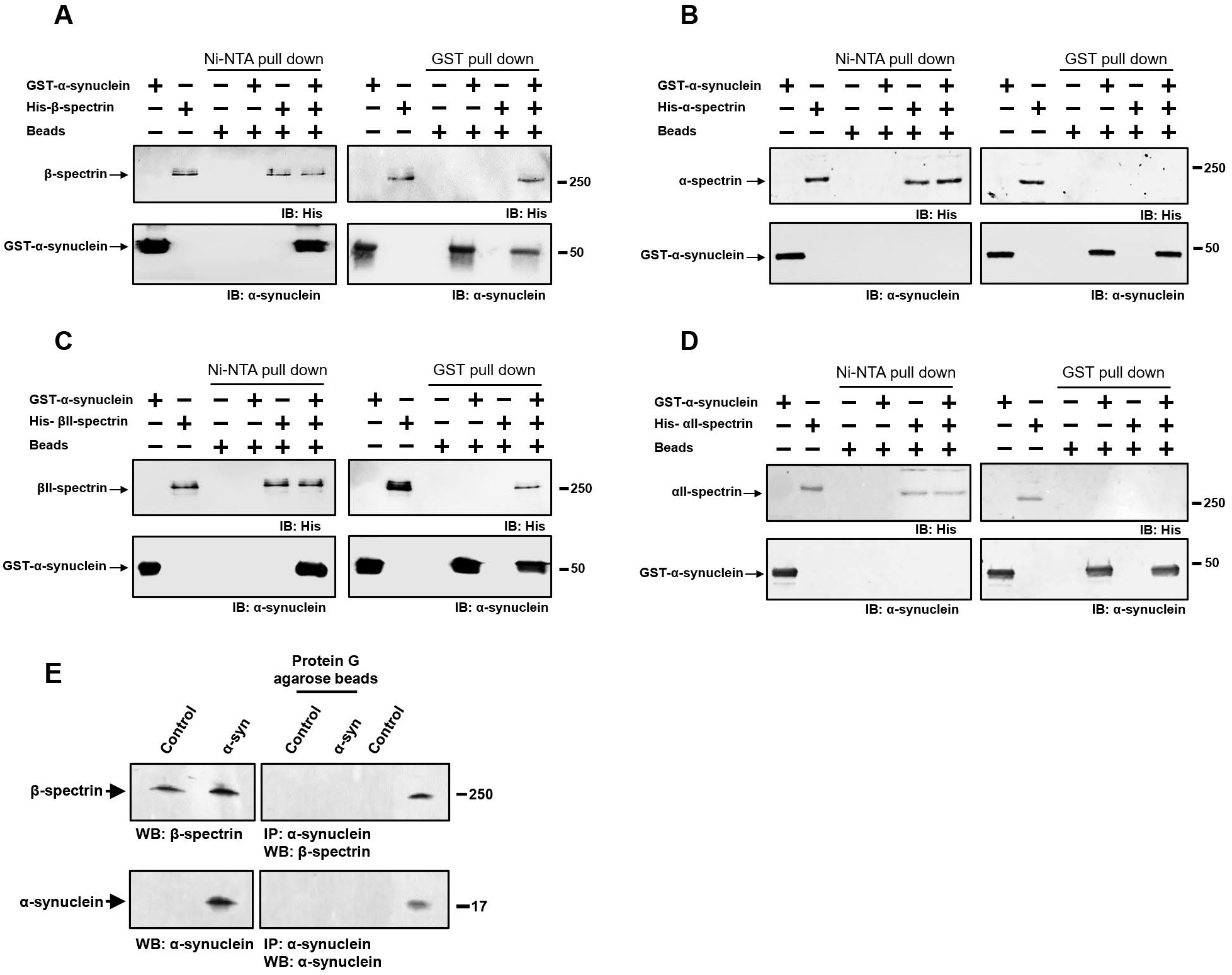
α-synuclein interacts directly with β-spectrin but not α-spectrin. *A*, Human α-synuclein and *Drosophila* β-spectrin interact in Ni-NTA and glutathione-S-transferase (GST) pull-down assays as monitored by immunoblotting for β-spectrin (His) or α-synuclein. *B*, No interaction of human α-synuclein and *Drosophila* α-spectrin in Ni-NTA or GST pull-down assays as monitored by immunoblotting for β-spectrin (His) or α-synuclein. *C*, Human α-synuclein and human βII-spectrin interact in Ni-NTA and GST pull-down assays as monitored by immunoblotting for βII-spectrin (His) or α-synuclein. *D*, No interaction of human α-synuclein and human αII-spectrin in Ni-NTA or GST pull-down assays as monitored by immunoblotting for αII-spectrin (His) or α-synuclein. *E*, Immunoprecipitation of α-synuclein in control and α-synuclein transgenic flies shows an association between α-synuclein and β-spectrin. Flies are 10 days old.

β-spectrin is a modular protein with multiple domains mediating interactions with specific partners in the cell (Fig. 4*A*). We have previously described β-spectrin transgenic flies expressing β-spectrin lacking ankyrin-binding activity (β-spec^Δank^) or β-spectrin with deletion of the pleckstrin homology (PH) domain (β-spec^ΔPH^) at levels similar to the endogenous protein and with similar levels of mutant compared to wild type transgenic β-spectrin (Fig. 4*A*, Extended Data Fig. 1-1) (Das et al., 2006). In the case of β-spec^ΔPH^ a nonsense mutation was introduced immediately upstream of the PH domain, resulting in expression of a slightly truncated protein. In β-spec^Δank^ an entire spectrin repeat (repeat 15) containing the ankyrin binding site was excised from β-spectrin and replaced with a repeat from α-spectrin (repeat 12) that preserves the structure of β-spectrin, but removes ankyrin binding activity. Expression of β-spec^ΔPH^ rescued locomotor deficits (Fig. 4*B*) and neurodegeneration (Fig. 4*C-F*) in α-synuclein transgenic flies. In contrast, expression of β-spec^Δank^ did not rescue locomotor defects (Fig. 4*B*) or neurodegeneration (Fig. 4*C-F*), implicating ankyrin binding in α-synuclein neurotoxicity.

**Figure 4.**
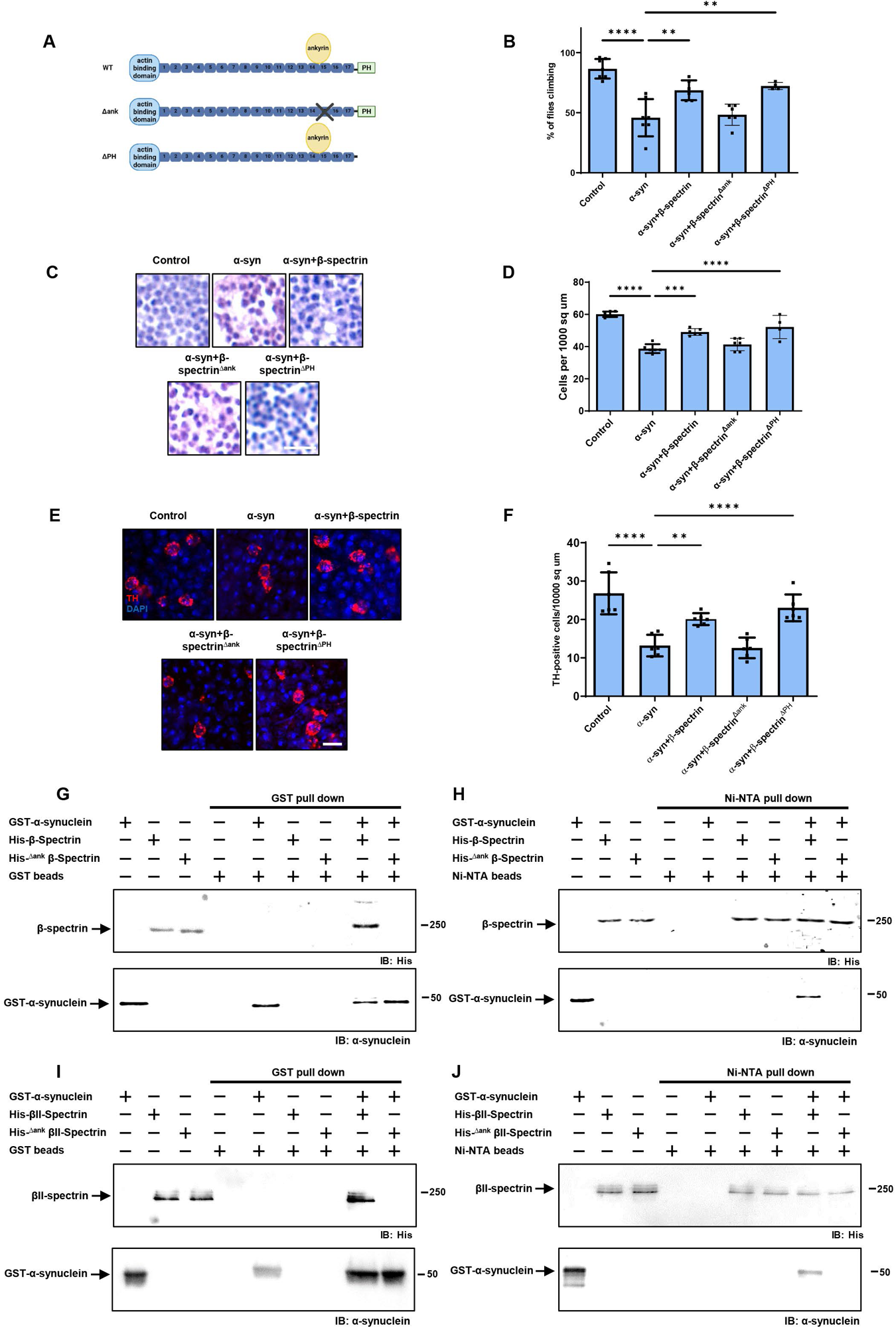
The ankyrin binding domain of β-spectrin mediates interactions with α-synuclein. *A*, Schematic diagram of the domains of wild type β-spectrin and the two mutant versions of spectrin used (β-spec^Δank^ and β-spec^ΔPH^). *B*, Climbing activity in control and human α-synuclein transgenic *Drosophila* with and without elevated expression of wild type and mutant forms of β-spectrin. n = minimum of 60 flies per genotype (six biological replicates of 10 flies each). *C,D*, Brain degeneration assayed by hematoxylin staining in the anterior medulla of control and α-synuclein transgenic *Drosophila* with and without elevated expression wild type and mutant forms of β-spectrin. n=6. *E,F*, Immunostaining for tyrosine hydroxylase in the anterior medulla of control and α-synuclein transgenic *Drosophila* with and without elevated expression wild type and mutant forms of β-spectrin. n=6. *G,H*, Human α-synuclein and wild type *Drosophila* β-spectrin but not β-spec^Δank^ interact in GST (*G*) or Ni-NTA (*H*) pull-down assays as monitored by immunoblotting for β-spectrin (His) or α-synuclein. *I,J,* Human α-synuclein and wild type βII-spectrin but not βII-spec^Δank^ interact in GST (*I*) or Ni-NTA (*J*) pull-down assays as monitored by immunoblotting for β-spectrin (His) or α-synuclein. All flies are 10 days old. Control flies are *nSyb-QF2, nSyb-GAL4/+.* The scale bars represent 5 µm (*A*) and 10 µm (*C*). Data are presented as mean ± SEM; P values determined with one-way ANOVA with Bonferroni post hoc test.

We next purified bacterially expressed, histidine-tagged β-spec^Δank^ and assessed binding to GST-tagged human α-synuclein. In contrast to wild type β-spectrin, β-spec^Δank^ failed to associate with α-synuclein as assayed by either GST (Fig. 4*G*) or Ni-NTA precipitation (Fig. 4*H*). We created a mutant in human βII-spectrin similar to fly β-spec^Δank^ by replacing the human βII-spectrin repeat 15 with the human α-spectrin repeat 12. We expressed the histidine-tagged mutant human protein in bacteria, purified the protein, and assessed binding to GST-tagged human α-synuclein. As observed with the *Drosophila* β-spec^Δank^ protein, the human βII-spec^Δank^ failed to co-precipitate with GST-tagged human α-synuclein (Fig. 4*I,J*). These biochemical data demonstrate that β-spectrin repeat 15 is critical for binding of α-synuclein to β-spectrin as well as for neurotoxicity (Fig. 4*B-F*).

Spectrin is present at the inner surface of the plasma membrane in a complex including ankyrin, the Na^+^/K^+^ ATPase and cell adhesion molecules (Das et al., 2006; Dubreuil, 2006; Mazock et al., 2010; Morrow and Stankewich, 2021; Teliska and Rasband, 2021). We examined the localization of ankyrin and Na^+^/K^+^ ATPase in α-synuclein transgenic flies by stimulated emission depletion microscopy (STED) following immunostaining of whole mount brains. In control flies the pattern of ankyrin overlapped that of β-spectrin, both localizing near the plasma membrane (Fig. 5*A-C*). The distribution of ankyrin was significantly altered in the α-synuclein transgenic flies, with an increase in cytosolic staining (Fig. 5*A,B*). Elevated expression of β-spectrin significantly restored the wild type distribution of ankyrin (Fig. 5*A-C*). Similarly, Na^+^/K^+^ ATPase redistributed to the cytoplasm from the plasma membrane upon expression of human α-synuclein in transgenic flies as monitored by immunostaining with the monoclonal antibody Nrv5F7, which recognizes the β subunit of the enzyme (Sun and Salvaterra, 1995). Expression of β-spectrin normalized the distribution of Na^+^/K^+^ ATPase (Fig. 5*D-F*).

**Figure 5.**
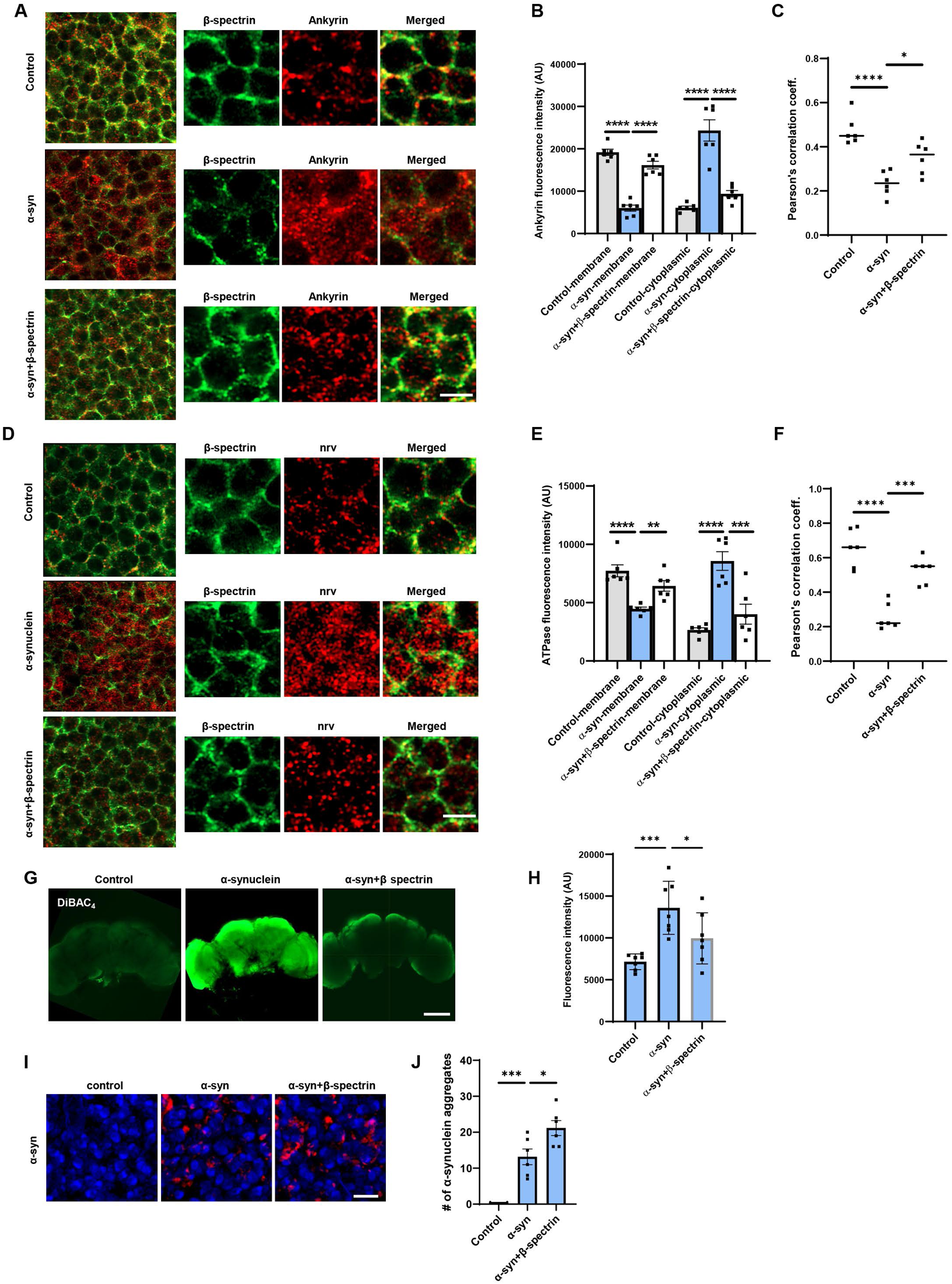
Expression of α-synuclein leads to mislocalization of ankyrin and Na^+^/K^+^ ATPase in *Drosophila* neurons. *A,B*, STED microscopy reveals colocalization of ankyrin (red) with subplasmalemmal β-spectrin (green) in wild type animals and increased cytoplasmic ankyrin (arrows) with expression of human α-synuclein. Elevated expression of β-spectrin partially normalizes ankyrin localization. n=6. *C*, Pearson’s correlation coefficient revealing an infrequent association between β-spectrin and ankyrin in α-synuclein transgenic *Drosophila.* Association is partially restored with elevated expression of β-spectrin. *D,E*, STED microscopy reveals substantial colocalization of Na^+^/K^+^ ATPase (red) with β-spectrin (green) in wild type animals and increased cytoplasmic Na^+^/K^+^ ATPase (arrows) with expression of human α-synuclein. Elevated expression of β-spectrin partially normalizes Na^+^/K^+^ ATPase localization. n=6. *F*, Pearson’s correlation coefficient revealing an infrequent association between β-spectrin and Na^+^/K^+^ ATPase in α-synuclein transgenic *Drosophila.* Association is partially restored with elevated expression of β-spectrin. *G,H*, Loss of plasma membrane polarization as monitored by DiBAC4(3) following expression of human α-synuclein. Elevated expression of β-spectrin partially normalizes membrane polarization. n=6. *I,J*, Immunofluorescence microscopy (*I*) and quantification (*J*) showing increased numbers of α-synuclein aggregates in neurons from fly brain sections of α-synuclein transgenic *Drosophila* with elevated β-spectrin. n=6. All flies are 10 days old. Controls are *nSyb, nSyb-GAL4/+.* The scale bars represent 2 µm (*A,C*), 5 µm (*I*) and 50 µm (*E*). Data are presented as mean ± SEM; P values determined with one-way ANOVA with Bonferroni post hoc test.

Na^+^/K^+^ ATPase plays a critical role in maintaining the ion gradients between the extracellular and intracellular environments, which in turn controls the resting membrane potential (Clausen et al., 2017). We assessed the effect of α-synuclein expression on membrane potential in flies by incubating dissected brains with the voltage-sensitive fluorescent dye bis-(1,3-dibutylbarbituric acid)-trimethine oxonol (DiBAC4(3)) (Bhavsar et al., 2019; Weiß and Bohrmann, 2019). α-synuclein transgenic flies showed significant depolarization in whole dissected brains in comparison to control flies, demonstrated as increased fluorescent intensity (Fig. 5*G,H*). Elevated expression of β-spectrin partially normalized membrane potential in whole mount brains from α-synuclein transgenic flies (Fig. 5*G,H*).

Aggregation of α-synuclein has been linked to neurotoxicity (Chen and Feany, 2005; Periquet et al., 2007; Lo Bianco et al., 2008; Shulman et al., 2011; Burré et al., 2018). We therefore assessed the number of aggregates present in α-synuclein transgenic flies with elevated levels of ß-spectrin. We found increased numbers of inclusions in ß-spectrin transgenic flies (Fig. 5*I,J*), consistent with our prior findings in α-spectrin transgenic animals (Ordonez et al., 2018).

To determine if results in the *Drosophila* α-synucleinopathy model extended to human cells, we used neurons differentiated from patient-derived induced pluripotent stem cells (iPSC) with a triplication of the α-synuclein locus and isogenic control cells (Devine et al., 2011; Ho et al., 2021). Immunostaining for βII-spectrin revealed the characteristic subplasmalemmal staining pattern in control neurons (Fig. 6*A*). In contrast, βII-spectrin immunostaining did not consistently show subplasmalemmal distribution in triplication neurons (Fig. 6*A*, arrows, *B*). We confirmed that human βII-spectrin co-immunoprecipitated with α-synuclein in homogenates from human neurons (Fig. 6*C*). We next examined ankyrin in human neurons. There are three mammalian ankyrin isoforms, ankryin-B, ankyrin-R and ankyrin-G. Ankryin-B is expressed in a widespread pattern in the nervous system, while ankyrin-R and ankyrin-G have more specific cellular and subcellular localization (Kordeli and Bennett, 1991; Lorenzo, 2020). When we stained for ankryin-B we observed increased cytoplasmic staining compared to isogenic controls (Fig. 6*D*, arrows, *E*), similar to findings in α-synucleinopathy model fly brains (Fig. 5*A,B*). We next used an antibody recognizing the βI subunit of human Na^+^/K^+^ ATPase to examine localization of Na^+^/K^+^ ATPase. As in α-synuclein transgenic *Drosophila* brains (Fig. 5*D,E*), we found increased cytosolic staining for human Na^+^/K^+^ ATPase in α-synuclein triplication neurons (Fig. 6*F*, arrows, *G*). Alteration of Na^+^/K^+^ ATPase localization was accompanied by plasma membrane depolarization as monitored by DiBAC4(3) fluorescence in human triplication neurons compare to isogenic controls (Fig. 6*H*).

**Figure 6.**
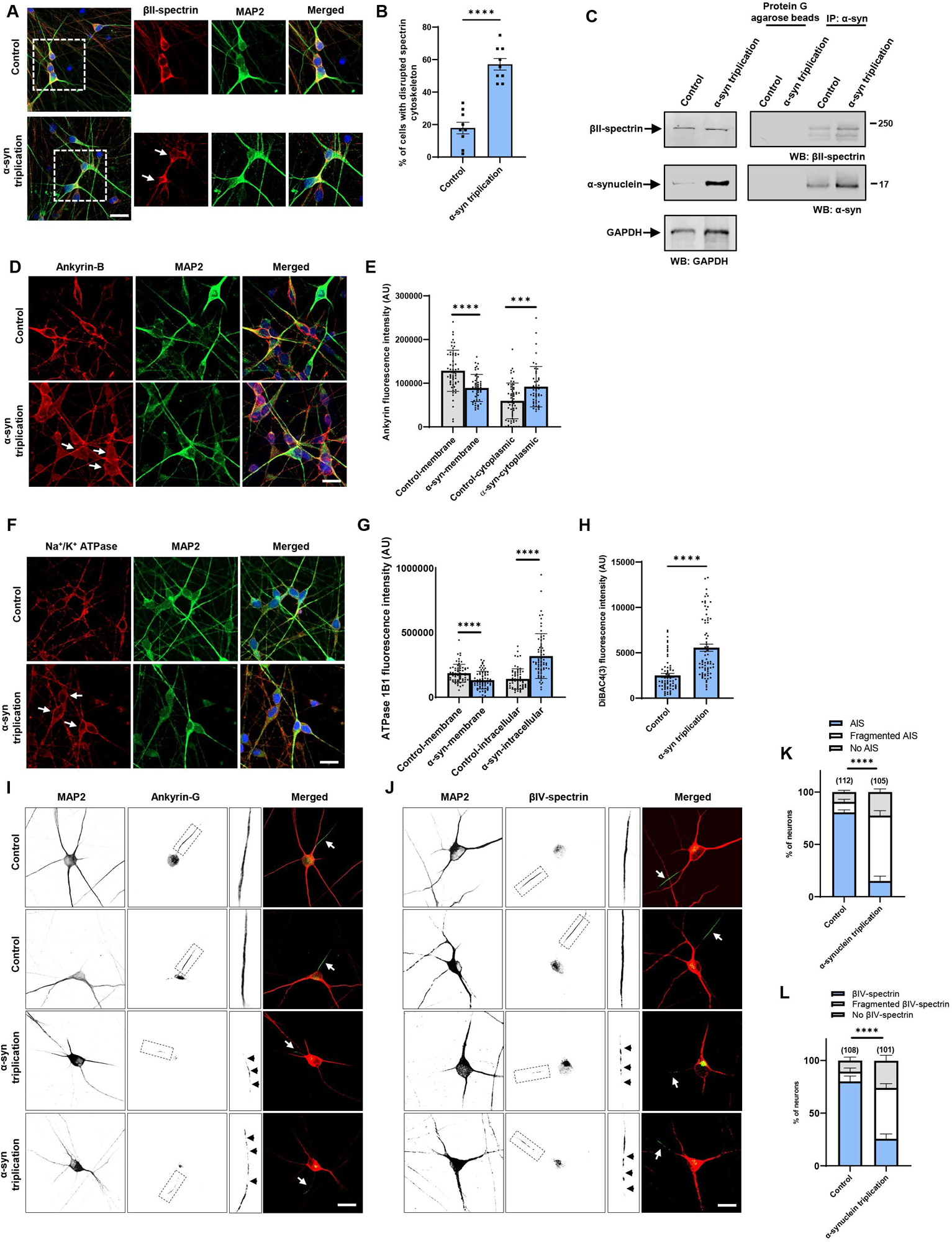
Increased expression of α-synuclein leads to disruption of the spectrin cytosketon and mislocalization of ankyrin and Na^+^/K^+^ ATPase in human neurons. *A,B*, The normal βII-spectrin immunoreactive subplasmalemmal network (red) is disrupted by increased expression of human α-synuclein in MAP2-positive (green) α-synuclein triplication patient neurons compared to isogenic control neurons. Arrows indicate cells with reduced βII-spectrin staining. n=9. C, Immunoprecipitation of α-synuclein in control and α-synuclein triplication patient derived neurons shows an association between α-synuclein and βII-spectrin. The blot is reprobed with an antibody to GAPDH to illustrate equivalent protein levels. *D,E,* Immunofluorescence microscopy reveals subplasmalemmal staining pattern for ankyrin-B (red) in MAP2-positive (green) control neurons and increased cytoplasmic staining in patient neurons (arrows). *F,G,* Immunostaining for Na^+^/K^+^ ATPase (red) shows a plasma membrane staining pattern in MAP2-positive (green) control neurons increased cytoplasmic staining in α-synuclein triplication patient neurons (arrows). *H*, Loss of plasma membrane polarization as monitored by DiBAC4(3) in α-synuclein triplication patient neurons compared to controls. *B,E,G,H,* Data are presented as mean ± SEM; P values determined with two-tailed t-test. *I-L*, Immunostaining for ankyrin-G (*I*) or ßIV-spectrin (*J*) identifies intact axon initial segments in most MAP2-positive control neurons (arrows) and increased numbers of neurons with loss or fragmentation of the axon initial segment (arrowheads) in α-synuclein triplication patient neurons, as quantified in (*K,L*). Assays were performed at 21 DIV. The scale bars represent 25 µm. Data are presented as mean ± SEM; P values determined with two-way ANOVA with Bonferroni post hoc test.

In addition to anchoring plasma membrane proteins such as Na^+^/K^+^ ATPase, ankyrins also organize discrete membrane domains of neurons. In particular, ankyrin-G localizes to and organizes the axon initial segment (Leterrier, 2018). We thus examined the axon initial segment in triplication and control neurons. Immunofluorescence using an antibody to ankyrin-G, identified a well-defined axon initial segment in most control neurons (Fig. 6*I*, arrows, *K*). In contrast, many triplication neurons displayed patchy staining for ankyrin-G, a pattern consistent with a fragmented axon initial segment (Galiano et al., 2012; Torii et al., 2020) (Fig. 6*I*, insets, arrowheads, *K*). We confirmed our findings with a second marker of the axon initial segment, ßIV-spectrin. Similar to ankyrin-G, immunostaining for ßIV-spectrin revealed decreased numbers of intact axon initial segments in triplication neurons (Fig. 6*J*, arrows, *L)*, with fragmentation of many axon initial segments (Fig. 6*J*, insets, arrowheads, *L*).

## Discussion

Here we demonstrate that pathological expression of α-synuclein in fly or human neurons leads to disruption of the subplasmalemmal spectrin network, mislocalization of ankyrins and Na^+^/K^+^ ATPase, consequent dysregulation of neuronal membrane potential and ultimately neurodegeneration. Our biochemical results suggest that disruption of the spectrin network reflects direct binding of α-synuclein to β-spectrin (Fig. 3). These findings are consistent with prior studies showing that α-synuclein colocalizes with β-spectrin in Lewy bodies from brains of patients with Lewy bodies (Leverenz et al., 2007). Intriguingly, β-spectrin has been linked genetically to the α-synucleinopathy dementia with Lewy bodies through genome association studies (Peuralinna et al., 2015). More generally, human genetic and animal model studies have demonstrated a critical role for spectrins in the development and function of the nervous system (Morrow and Stankewich, 2021). Mutations in the genes encoding spectin isoforms give rise to neurodegenerative ataxias (βIII-spectrin), neurodevelopmental and behavioral deficits (βII and βIII-spectrin) and deafness (βIV- and βV-spectrin) (Cousin et al., 2021; Morrow and Stankewich, 2021; Teliska and Rasband, 2021). Our current results suggest that the α-synucleinopathies may represent additional members of the human spectrinopathy family of neurological diseases.

Experiments in animal and cell culture models of α-synucleinopathy have implicated alterations in multiple fundamental cellular processes, including vesicular trafficking (Chung et al., 2013; Burré et al., 2018; Vidyadhara et al., 2019) mitochondrial dynamics and function (Martin et al., 2006; Ordonez et al., 2018; Sarkar et al., 2020; Portz and Lee, 2021), nuclear regulation (Kontopoulos et al., 2006; Pinho et al., 2019; Schaser et al., 2019; Vasquez et al., 2020) and proteostasis (Auluck et al., 2002; Colla et al., 2012; Yan et al., 2019; Karim et al., 2020; Sarkar et al., 2021) in neurotoxicity. Interestingly, spectrin has been implicated in the control of each of these cellular functions as well. Spectrin was originally identified as a key component of the subplasmalemmal cytoskeleton of the red blood cell critical to maintaining cell shape and integrity during mechanical deformation (Liem, 2016). Much subsequent investigation has focused on conceptually similar subplasmalemmal roles in a variety of cell types, particularly those, like cardiomyocytes, subject to mechanical stress. However, spectrin has also been localized to a variety of intracellular compartments, including transport vesicles, mitochondria and the nucleus (Zagon et al., 1986), where the protein complex has been implicated in vesicle transport and DNA damage repair, among other functions (Lambert, 2018, 2019; Goodman et al., 2019; Morrow and Stankewich, 2021). Thus, binding of α-synuclein to spectrin at multiple intracellular sites may perturb a number of key cell biological roles subserved by spectrin. Alternatively, we have previously described altered actin dynamics downstream of spectrin in α-synucleinopathy models, which controls mitochondrial dynamics and function. Some or all of the effects of α-synuclein may thus reflect altered organization and dynamics of the actin cytoskeleton following loss of β-spectrin binding to ankyrin and disruption of the subplasmalemmal spectrin network. Additional work will be needed to distinguish these possibilities.

In addition to the more general functions of spectrin in cellular biology, in neurons distinct spectrin isoforms organize and maintain specific subcellular domains, including the axonal initial segment and nodes of Ranvier, which are needed for initiation and propagation of action potentials (Teliska and Rasband, 2021). We show here that the axon initial segment is abnormal in neurons from patients with Parkinson’s disease due to triplication of the α-synuclein locus compared to isogenic control neurons (Fig. 6*I,J*). Spectrins also form a key component of the periodic rings of actin and spectrin known as membrane-associated periodic cytoskeleton present in axons (Xu et al., 2013) and dendrites (D’Este et al., 2015; Han et al., 2017). In mouse models disruption of either the membrane-associated periodic cytoskeleton upon loss of αII-spectrin (Huang et al., 2017) or nodes of Ranvier with genetic deletion of βI- and βIV-spectrin (Liu et al., 2020) leads to axonal degeneration. Alteration of specific neuronal structures and function by pathological binding of α-synuclein to β-spectrin may therefore promote neurotoxicity in α-synucleinopathy.

As well as colocalizing with α-synuclein in Lewy bodies, β-spectrin has also been reported to interact with α-synuclein in mouse brain (McFarland et al., 2008) and cultured cortical neurons (Chung et al., 2017). These findings raise the possibility that binding of α-synuclein to β-spectrin may be relevant to the normal function of α-synuclein. Although the precise role that α-synuclein plays in nervous system function remains unclear, multiple studies have demonstrated modulation of neurotransmitter release by α-synuclein, consistent with the presynaptic localization of the protein (Runwal and Edwards, 2021). Spectrins and ankyrins are also present in the presynapse (Zagon et al., 1986; Pielage et al., 2008; Smith et al., 2014) and might transduce or modulate the synaptic activity of α-synuclein.

The precise form of α-synuclein that interacts with β-spectrin in vivo requires further definition. In vitro (McFarland et al., 2008) and in vivo (Ordonez et al., 2018) evidence suggests that serine 129 phosphorylation promotes the interaction of α-synuclein with spectrin. Phosphorylation might influence the interaction of α-synuclein with β-spectrin directly, or might work through indirect effects on α-synuclein aggregation (Fujiwara et al., 2002; Ghanem et al., 2022). We and others have implicated oligomeric forms of α-synuclein in neurotoxicity, with evidence for a protective role for larger inclusions, perhaps as a sink for toxic smaller aggregates (Chen and Feany, 2005; Periquet et al., 2007; Chen et al., 2009; Olsen and Feany, 2021; Panicker et al., 2021). Our current observation that large inclusions increase in numbers when α-synuclein neurotoxicity is reduced by elevating β-spectrin levels (Fig. 5*I,J*) is consistent with these prior data.

We show here that levels of β-spectrin strongly influence the ability of human α-synuclein to show toxicity to dopaminergic and non-dopaminergic neurons in vivo. In particular, we find that increasing β-spectrin can protect from α-synuclein neurotoxicity by restoring the normal organization of the subplasmalemmal spectrin network and maintaining normal localization and activity of Na^+^/K^+^ ATPase (Figs. 1,5). These results suggest that therapeutic strategies aimed at stabilization of the spectrin cytoskeleton (Morrow and Stankewich, 2021) or normalization of the activity of downstream targets represent potential new approaches to the treatment of Parkinson’s disease and related α-synucleinopathies.

## Acknowledgements

Fly stocks obtained from the Bloomington *Drosophila* Stock Center (NIH P40OD018537) and C. Potter were used in this study. Monoclonal antibodies were obtained from the Developmental Studies Hybridoma Bank developed under the auspices of the NICHD and maintained by the University of Iowa, Department of Biology, Iowa City, IA 52242, and the UC Davis/NIH NeuroMab Facility. Dr. M. Rasband kindly provided the ßIV-spectrin antibody. This work was supported by NIH-NINDS R01NS098821. This research was funded in part by Aligning Science Across Parkinson’s [ASAP-000301] through the Michael J. Fox Foundation for Parkinson’s Research (MJFF). For the purpose of open access, the author has applied a CC BY public copyright license to all Author Accepted Manuscripts arising from this submission.

---

**Extended Data Figure 1-1.**
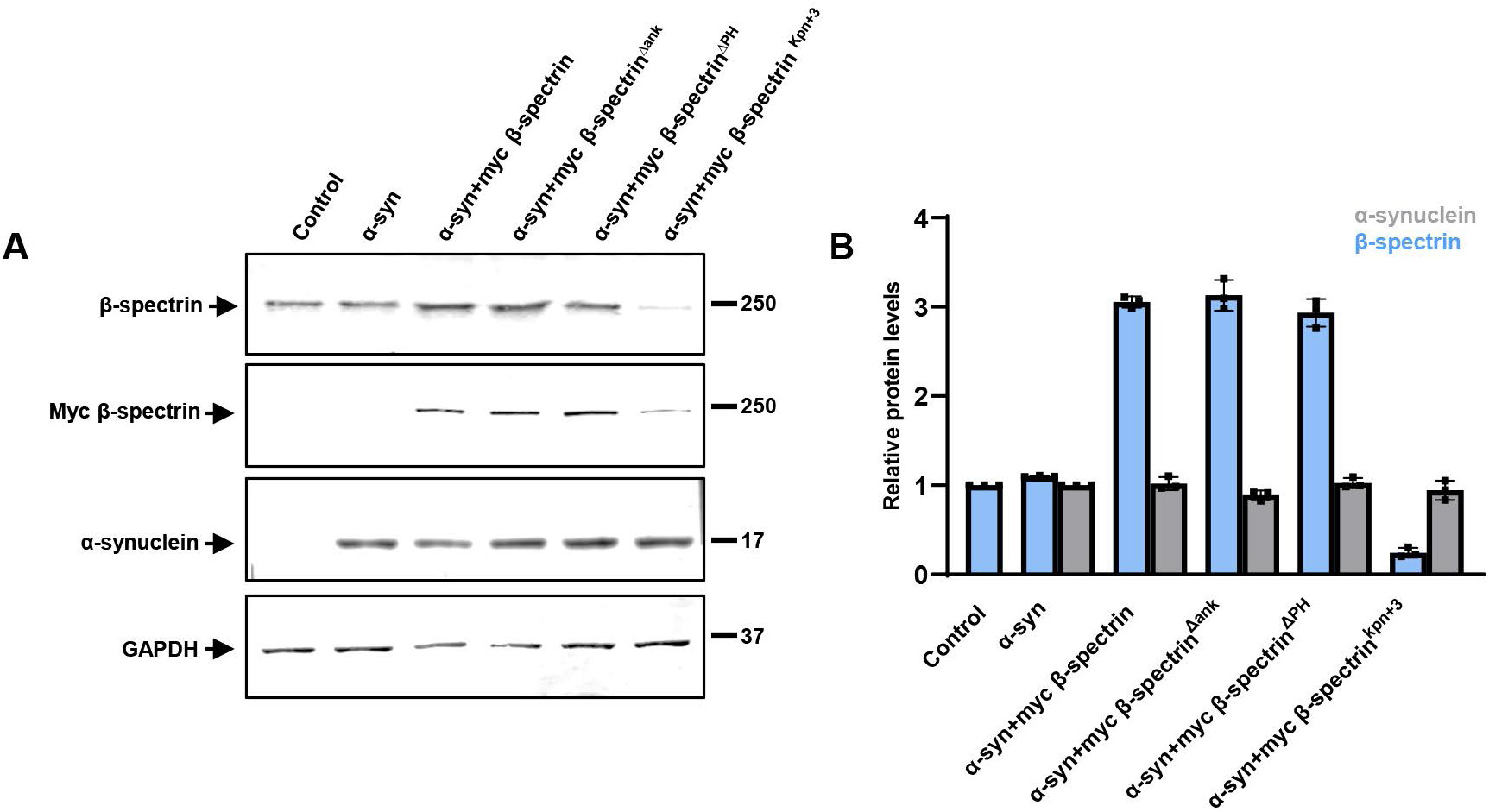
β-spectrin and α-synuclein expression in transgenic *Drosophila* heads. *A,B*, Immunoblotting analysis using an antibody to the myc tag present on transgenic β-spectrin reveals equivalent levels of expression of wild type β-spectrin, β-spectrin^Δank^ and β-spectrin^ΔPH^, and reduced levels of the β-spectrin^Kpn+3^ variant. The blot is reprobed with an antibody to GAPDH to illustrate equivalent protein levels. Relative β-spectrin protein levels are normalized to control (*nSyb-GAL4, nSyb-QF2/+*). Relative α-synuclein protein levels are normalized to α-synuclein (*QUAS-alpha-synuclein, nSyb-GAL4, nSyb-QF2/+*). n=3. Flies are 1 day old.

